# Investigating crosstalk among PTMs provides novel insight into the structural basis underlying the differential effects of Nt17 PTMs on mutant Httex1 aggregation

**DOI:** 10.1101/2021.02.21.432155

**Authors:** Anass Chiki, Zhidian Zhang, Kolla Rajasekhar, Luciano A. Abriata, Iman Rostami, Lucien Krapp, Driss Boudeffa, Matteo Dal Peraro, Hilal A. Lashuel

**Author notes:** These authors contributed equally to this work (Anass Chiki, Zhidian Zhang).

## Abstract

Post-translational modifications (PTMs) within the first 17 amino acids (Nt17) of the Huntingtin protein (Htt) have been shown to inhibit the aggregation and attenuate the toxicity of mutant Htt proteins *in vitro* and in various models of Huntington’s disease. Our group’s previous studies suggested that the Nt17 PTM code is a combinatorial code that involves a complex interplay between different PTMs. Here, we expand on these studies by investigating the effect of methionine 8 oxidation (oxM8) and crosstalk between this PTM and either lysine 6 acetylation (AcK6) or threonine 3 phosphorylation (pT3) on the aggregation of mutant Httex1. We show that M8 oxidation delays but does not inhibit the aggregation and has no effect on the final morphologies of mutant Httex1 aggregates. This delay in aggregation kinetics could be attributed to the transient accumulation of oligomeric aggregates, which disappear upon the formation of Httex1 oxM8 fibrils. Interestingly, the presence of both oxM8 and AcK6 resulted in dramatic inhibition of Httex1 fibrillization, whereas the presence of oxM8 did not influence the aggregation inhibitory effect of pT3. To gain insight into the structural basis underlying these proteins’ aggregation properties, we investigated the impact of each PTM and the combination of these PTMs on the conformational properties of the Nt17 peptide by circular dichroism spectroscopy and molecular dynamics simulation. These studies show that M8 oxidation decreases the helicity of the Nt17 in the presence or absence of PTMs and provides novel insight into the structural basis underlying the effects of different PTMs on mutant Httex1 aggregation. PTMs that lower the mutant Httex1 aggregation rate (oxM8, AcK6/oxM8, pT3, pT3/oxM8, and phosphorylation at Serine 13) result in stabilization and increased population of a short N-terminal helix (first eight residues) in Nt17 or decreased abundance of other helical forms, including long helix and short C-terminal helix. PTMs that did not alter the aggregation of mutant Httex1 exhibit a similar distribution of helical conformation as the unmodified peptides. These results show that the relative abundance of N- vs. C-terminal helical conformations and long helices, rather than the overall helicity of Nt17, better explains the effect of different Nt17 PTMs on mutant Httex1; thus, explaining the lack of correlation between the effect of PTMs on the overall helicity of Nt17 and mutant Httex1 aggregation *in vitro*. Taken together, our results provide novel structural insight into the differential effects of single PTMs and crosstalk between different PTMs in regulating mutant Httex1 aggregation.

**TOC Figure:** 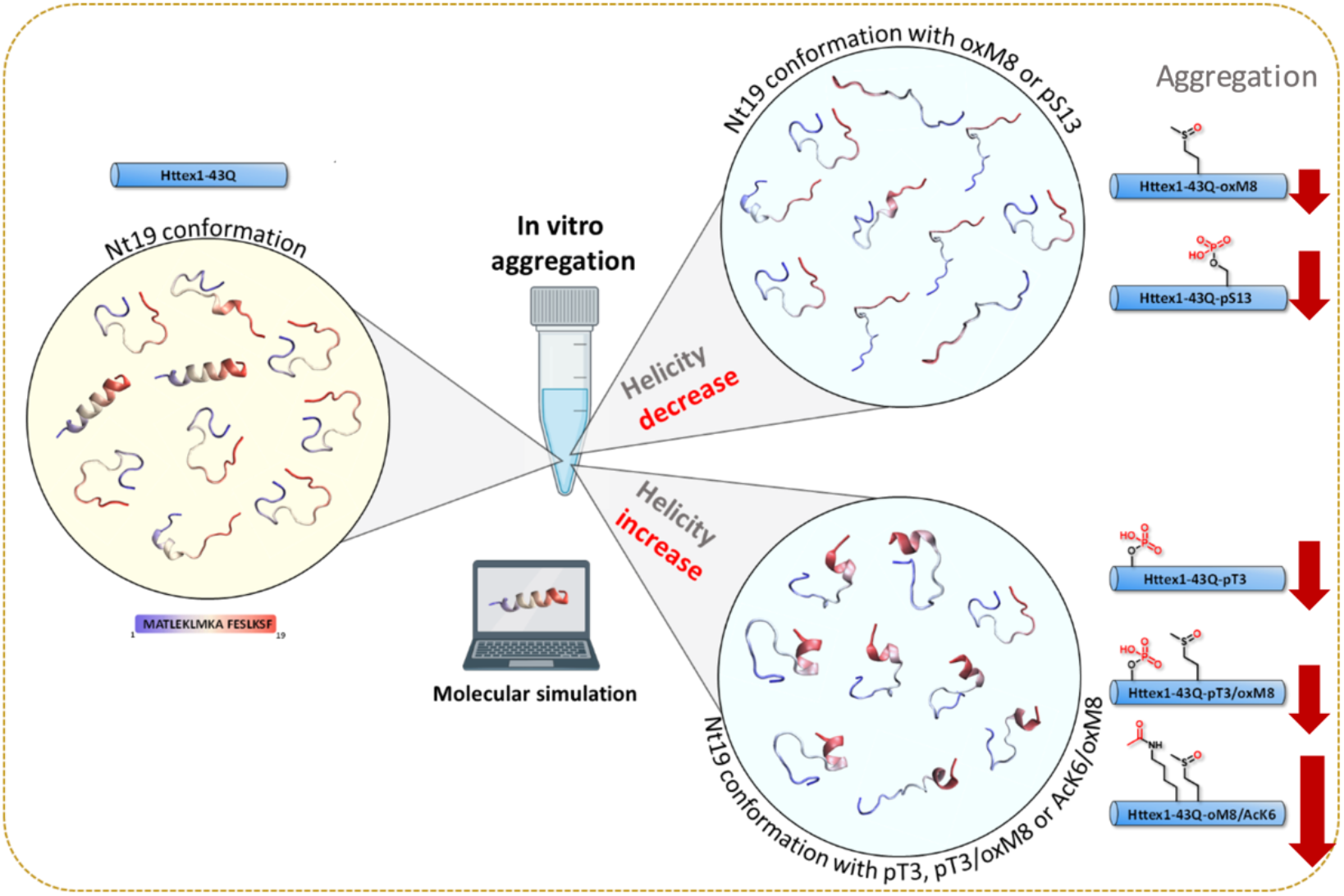

## Introduction

Huntington’s disease (HD) is a fatal, autosomal neurodegenerative disease characterized by motor [1, 2] and cognitive declines [3] as well as psychiatric symptoms [4]. HD is caused by a mutation in the huntingtin gene (*HTT*), resulting in an expansion in the CAG repeat within its first exon [5, 6], which is then translated into an extended polyglutamine (polyQ) repeat in the huntingtin protein (Htt) [7]. HD occurs when the length of the polyQ repeat is higher than the critical threshold of ≥36 [8]. At the neuropathological level, HD is characterized by neuronal degeneration in the striatum and the cortex [9, 10] and the formation and accumulation of nuclear inclusions composed of mutant Htt (polQ repeat ≥36) aggregates and other proteins [11, 12]. Several studies have shown that these inclusions are composed of fibrillar and potentially oligomeric species derived from N-terminal fragments containing expanded polyQ repeats [11, 13–16]. One of the major N-terminal fragments found in these inclusions represents an N-terminal fragment that corresponds to exon1 of the Htt protein (Httex1) [14, 16]. Overexpression of mutant Httex1 alone with polyQ length ranging from 80 to 175 reproduces many aspects of HD pathology in various animal and cellular models, including the formation of huntingtin inclusions [13, 17–19].

Although increasing evidence suggests that mutant Htt aggregation and toxicity play central roles in HD’s pathogenesis, the molecular events responsible for triggering mutant Htt aggregation, the nature of the toxic species, and the mechanisms by which they cause neurodegeneration remain unknown. Initial efforts focused on disentangling the relationship between Htt aggregation and toxicity and HD have focused on identifying modifiers of Htt aggregation based on targeting the polyQ repeat domain, which has been shown to be the primary sequence responsible for initiating Htt aggregation. However, recent studies suggest that post-translation modifications (PTMs) in close proximity or far from the polyQ domain have the potential to modify not only mutant Htt levels and functions, but also its aggregation and toxicity [20–27]. Interestingly, several of these PTMs occur within the first N-terminal 17 amino acids of Htt, directly flanked by the polyQ domain (Figure 1-A). These modifications include phosphorylation at multiple serine and threonine residues (T3, S13, and S16), acetylation, ubiquitination, and SUMOylation at selected lysine residues (K6, K9, K15) (Figure 1-A). Mutating both S13 and S16 to aspartate to mimic the phosphorylation was shown to reverse the pathology of mutant Htt in an HD mouse model [28] and modulate Htt aggregation in different cellular models [13, 17–19]. Recently, we showed that phosphorylation at T3, S13, and/or S16 inhibited the aggregation of WT and mutant Httex1 *in vitro* [29, 30].

**Figure 1.**
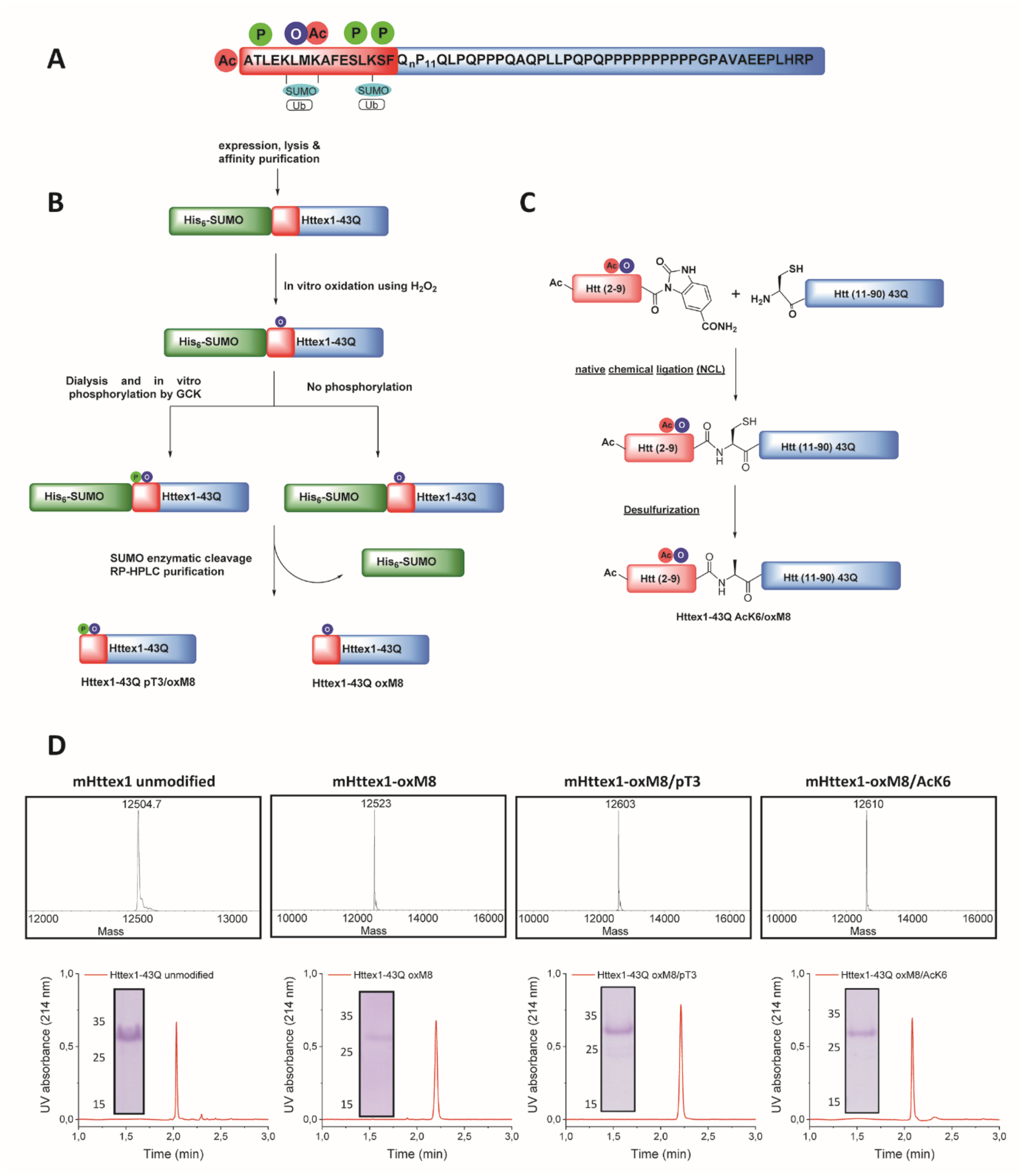
Chemical and enzymatic methods for the generation of of Httex1-43Q-oxM8, Httex1-43Q-oxM8/pT3, and Httex1-43Q-oM8/AcK6. **(A)** Schematic presentation of the Httex1 sequence, highlighting the cluster of PTMs in the Nt17 domain. **(B)** Schematic presentation of the SUMO-based strategy used to produce Httex1-43Q-oxM8 and Httex1-43Q-oxM8/pT3. The SUMO-Httex1-43Q was produced and purified by Nickel IMAC purification and subsequently oxidized or both oxidized and phosphorylated by GCK. Next, the SUMO tag was removed by ULP1, and the desired protein was purified by RP-HPLC. **(C)** Schematic representation for the semisynthetic strategy used for the generation of mHttex1-oxM8/AcK6 (adapted from [30]). **(D)** Characterization of Httex1-43Q-oxM8, Httex1-43Q-oxM8/pT3, Httex1-43Q-oM8/AcK6, and unmodified Httex1-43Q by ESI/MS, UPLC, and SDS-PAGE.

Furthermore, TBK1-mediated phosphorylation of mutant Httex1 at S13 and S16 lowers its levels and results in a significant reduction in Httex1 aggregation and inclusion formation in different cellular models and a *C. elegans* model of HD [31]. These findings, combined with our observation that the levels of phosphorylated Htt at T3 are decreased in pathological conditions [32], suggest that phosphorylation within the Nt17 domain protects against mutant Htt aggregation. Similarly, SUMOylation at multiple N-terminal lysine residues inhibits Httex1 aggregation *in vitro* [33]. Interestingly, although lysine acetylation at K6, K9, or K15 does not significantly alter the aggregation profile of Httex1 [30], acetylation at K6 was shown to significantly inverse the inhibitory effects of phosphorylation at T3 on aggregation [30]. Similarly, phosphorylation of S13 and S16 was shown to regulate Httex1 acetylation at K9 [20], and ubiquitination/SUMOylation at K6 and K9 [20, 27]. Finally, competing modifications, such as ubiquitination and SUMOylation, were shown to exert different effects on Htt levels and degradation [22]. Taken together, these findings, combined with the fact that multiple reversible PTMs cluster within eight amino acids in Nt17, suggest that many of these modifications act in concert rather than individually and that the Htt PTM code involves crosstalk between different PTMs, in particular, those that exist close to each other.

One PTM that remains unstudied and its effect on mutant Htt aggregation remains unknown is the oxidation of methionine 8 (oxM8). This residue is highly conserved [34], and the levels of M8 oxidation were shown to be higher in the R6/2 HD mouse model [35]. However, the absence of immunochemical methods to detect M8 oxidation has precluded studies aimed at understanding its relevance and potential roles in HD neuropathology. Despite this, methionine oxidation is one of the most common modifications that occur under oxidative stress [36], which has been linked to the pathogenesis of HD [37] and other neurodegenerative diseases. Elevated levels of oxidative stress markers, including reactive oxygen species (ROS), were found in blood [38] and postmortem brains of HD patients [39]. Although there is no direct evidence establishing that M8 is oxidized, several studies have suggested that this could occur under oxidative stress conditions and that M8 oxidation could act as a sensor of ROS to regulate Htt phosphorylation and localization [34, 40].

Mitomi and colleagues [35] showed, using hydrogen peroxide (H_2_O_2_) mediated oxidizing conditions *in vitro*, that oxidation at M8 occurs only post-aggregation on non-soluble forms of mutant Httex1. Recent studies based on quantitative NMR showed that TiO_2_ nanoparticles induced M8 oxidation and abolished the aggregation of a model peptide of Httex1, and reduced the model peptide binding to lipids micelles [41]. However, most of these studies were carried out either using Nt17 peptides, GST-tagged Httex1 protein [35], or a model peptide consisting of only the Nt17 linked to short polyQ tracts of only 7 or 10 glutamines, lacking the proline-rich domain and most the C-terminal domain of Httex1 [41]. Additionally, in many studies, the H_2_O_2_ was not removed from the solution prior to the aggregation experiments.

Here, we used an integrative approach combining biophysical and computational studies to gain insight into the effect of M8 oxidation and potential crosstalk between M8 oxidation with neighboring PTMs in regulating the structure and aggregation of mutant Httex1. Towards this goal, we used chemoenzymatic approaches to produce site-specific modified Httex1 proteins bearing different post-translationally modified proteins, including: 1) oxM8, 2) oxM8 and phosphorylation at T3, and 3) oxM8 and acetylation at K6. With these proteins in hand, we performed systematic studies to understand the effect of M8 oxidation and its crosstalk with pT3 and AcK6 in modulating the structure, aggregation, and fibrillar morphology of Httex1. To further understand the structural basis underlying the effects of these modifications on mutant Httex1 aggregation, we performed circular dichroism (CD) analysis and atomistic molecular simulations focusing on investigating how these different combinations of Nt17 PTMs influence Nt17 structure and helicity. Our findings provide new insight into the PTM code of Nt17, and the differential effects of PTM-induced conformational changes on mutant Httex1 (mHttex1) aggregation.

## Results

### Production of oxidized mutant Httex1 proteins

Httex1 contains two methionine residues within its Nt17 domain. The N-terminal methionine is cleaved by aminopeptidase resulting in N-terminally acetylated alanine as the first residue [42]. This means that once expressed, Httex1 contains only a single methionine residue at position 8 (M8). Investigating the biological role of M8 oxidation is more challenging than other Nt17 PTMs, such as phosphorylation, because there are no natural amino acids that mimic methionine oxidation and factors that are known to induce protein oxidation, such as oxidative stress, are not always chemoselective. Furthermore, it is not always possible to assess the extent of site-specific methionine oxidation under cellular conditions, e.g., oxidative stress, that is known to induce M8 oxidation. Therefore, to investigate the effect of M8 oxidation on Httex1 aggregation, we sought to prepare mutant Httex1 proteins that are homogeneously oxidized at M8. Several protocols have been used to oxidize methionine residues in proteins *in vitro*, including treatment with H_2_O_2_, transition metal ions, 2,2-azobis(2-amidinopropane) dihydrochloride (AAPH), tert-butyl hydroperoxide (t-BHP), and UV exposure [43–46].

In the context of Httex1, previous studies have shown that treatment with H_2_O_2_ allows for efficient oxidation of M8 within a GST-Httex1 fusion protein [35], the Nt17 [34], or Httex1 model peptide consisting of only the Nt17 domain with ten additional glutamine residues (Nt17Q10) [41]. Some of the limitations of these studies are: 1) none of the proteins used to represent the native sequence of Httex1; 2) the extent of oxM8 was always confirmed by mass spectrometry or other methods; or 3) that the structural or aggregation properties of the protein and peptide models was assessed in the presence of the oxidizing agent, H_2_O_2_. It is known that incubation of proteins under such harsh conditions for an extended period might modify other residues within the sequence, such as histidine or phenylalanine [47], which could alter the sequence and biophysical properties of Httex1.

To overcome these limitations, we developed a strategy for producing native mutant Httex1 specifically oxidized at M8 (oxM8). We took advantage of recent advances from our lab that allow for the generation of microgram quantities of highly pure mutant Httex1 fused to the SUMO protein [48], which enhances the solubility of mutant Httex1 (Figure 1-B). The desired PTMs (oxidation or phosphorylation) are then introduced, using enzymatic or chemical approaches, into Nt17 of the fusion protein, followed by removal of the SUMO protein and purification of the modified mutant Httex1 by RP-HPLC[49].

Mutant Httex1, with 43 glutamine residues (mHttex1) fused to a SUMO tag protein at its N-terminal (SUMO-mHttex1), was expressed and purified as previously described (Figure S1-A) [48]. Next, the fusion protein was subjected to oxidation using 400 mM of H_2_O_2_ in 50mM Tris, 500 mM NaCl, 500 mM Imidazole, pH 7.5. The oxidation reaction was monitored over time by ESI/MS (Figure S1-C). For each time point, an analytical cleavage of the SUMO tag using the ULP1 enzyme was performed, and the resulting cleaved product was analyzed by ESI/MS. As shown in Figure S1-C, the M8 residue was completely oxidized after 2 h of incubation with H_2_O_2_. The oxidized SUMO-mHttex1 was then subjected to ULP1 cleavage. Upon verifying the complete removal of the SUMO tag, m-Httex1-oxM8 was immediately purified by reverse phase HPLC (RP-HPLC) (Figure S1-E). The fractions containing the protein of interest were then pooled together and lyophilized. The final purity of the mHttex1-oxM8 was verified by ESI/MS, UPLC, and SDS-PAGE (Figure 1-D).

To investigate the effect of potential crosstalk between oxM8 and other Nt17 PTMs on mutant Httex1 aggregation, we generated mutant Httex1 proteins oxidized at M8 and phosphorylated at T3 (pT3) or acetylated at K6 (AcK6). To produce mHttex1 oxidized at M8 and phosphorylated at T3 (mHttex1-oxM8/pT3), we first generated SUMO-mHttex1-oxM8 as described above. After overnight dialysis to remove the excess of H_2_O_2_, the fusion protein was co-incubated overnight with GCK kinase, a kinase that we recently reported to efficiently phosphorylate Httex1 specifically at T3[49]. The extent of phosphorylation was monitored by ESI/MS. As shown in Figure S1-D, the SUMO-mHttex1-oxM8 showed an additional +80 Da, indicating the addition of a single phosphate group. The SUMO-mHttex1-oxM8/pT3 was then subjected to UPL1 cleavage followed by RP-HPLC purification of mHttex1-oxM8/pT3 (Figure S1-F). The protein’s purity was verified by ESI/MS, UPLC, and SDS-PAGE (Figure 1-D).

To produce mutant Httex1 that is both oxidized at M8 and acetylated at lysine 6, we employed a semisynthetic protein strategy (Figure 1-C) that we previously used to introduce single or multiple PTMs within Nt17 of mHttex1 [49]. mHttex1-AcK6/oxM8 was produced using native chemical ligation between the Htt A10C-90 43Q and Ac-2-9-Nbz AcK6/oxM8 peptides (characterization by ESI-MS and UPLC is shown in Figure S2-B), followed by desulfurization to convert the cysteine to the native Alanine 10 (Figure S2-C). The purity of mHttex1 AcK6/oxM8 was determined by SDS-PAGE, UPLC, and ESI/MS (Figure 1-D).

### Oxidation at M8 delays the aggregation of mutant Httex1

To determine the effect of oxM8 on mutant Httex1, mHttex1-oxM8 and mHttex1 were subjected to a disaggregation protocol, as previously reported [48]. Any remaining aggregates were removed by filtration through a 100 kDa filter before initiating the aggregation. The extent of and kinetics of aggregation were monitored by quantifying the amount of soluble proteins at different time points using a previously described UPLC-based sedimentation assay [50–52]. Also, the effect of oxidation on the secondary structure and morphology of the fibrils was also assessed by circular dichroism spectroscopy (CD) and electron microscopy (EM). Figure 2-A shows the percentage of the remaining soluble protein over time. As expected, unmodified mHttex1 exhibited almost complete depletion of the soluble monomer (Figure 2-A) and full conversion into aggregates after 48 h, which was confirmed by a shift from random to a β-sheet structure by CD (Figure 2-B) and the presence of mature long fibrils as discerned by EM (Figure 2-C). In contrast, mHttex1-oxM8 showed a delay in aggregation compared to the unmodified mHttex1 (Figure 2-A). The aggregation of mHttex1-oxM8 showed 70% and 29% remaining monomer after 12 and 24 h, respectively, compared to 38% and 7% for the unmodified mHttex1 (Figure 2-A). However, after 48 h, both proteins exhibited complete aggregation as discerned by the complete disappearance of soluble Httex1 (Figure 2-A) and the CD spectra of both proteins, which showed a signal that corresponds to predominantly β-sheet-rich structures (Figure 2-B). The fibrils formed by mHttex1 oxM8 after 48 h were similar to those formed by the unmodified mutant Httex1, suggesting that oxM8 influences aggregation’s kinetics but not the final structure of the fibrils (Figure 2-C).

**Figure 2.**
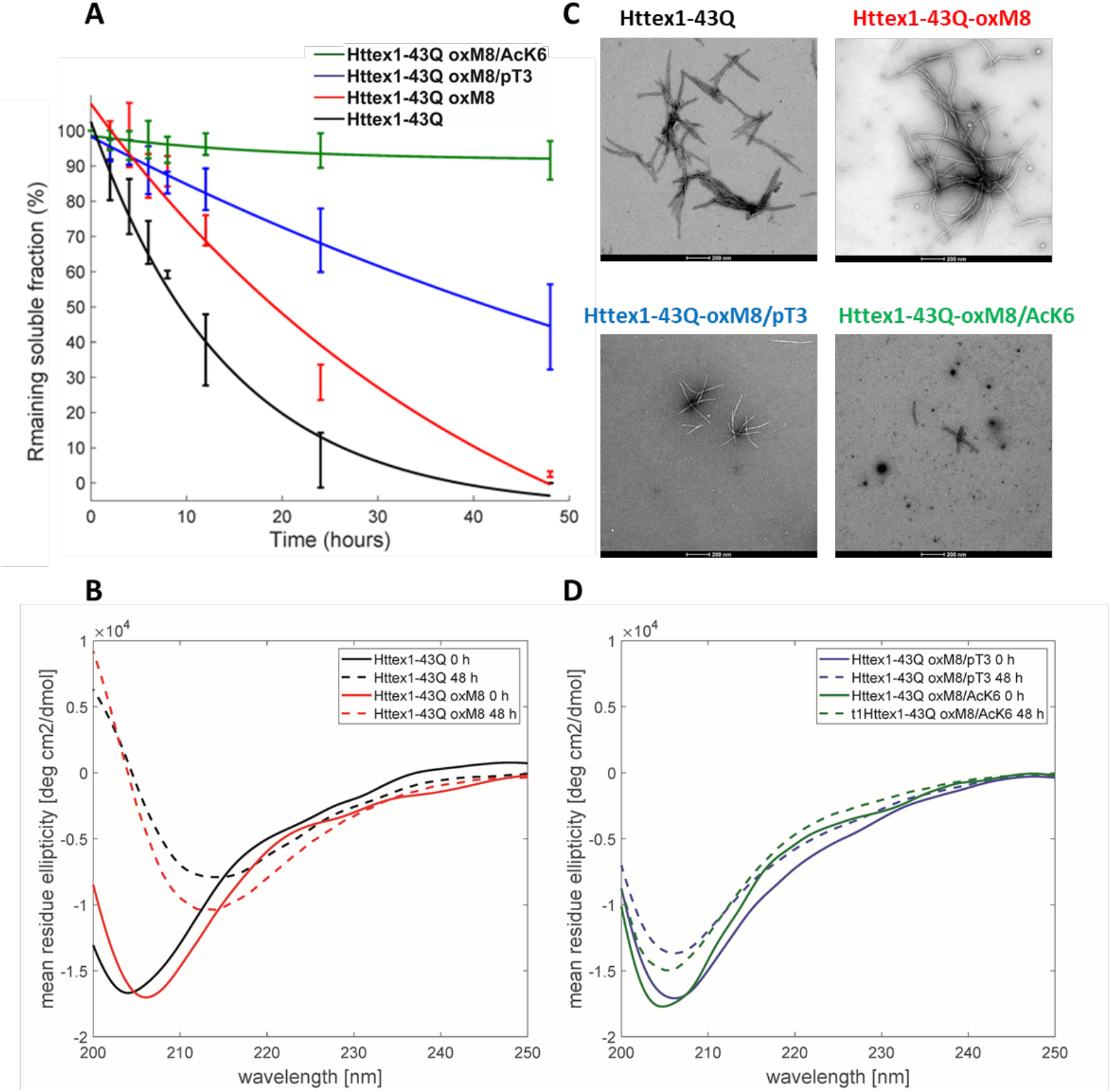
Biophysical studies to elucidate the effect of pT3, OxM8, and AcK6 on mutant Httex1 aggregation. **(A)** Httex1 aggregation studies (at 5 μM) of Httex1-43Q-oxM8, Httex1-43Q-oxM8/pT3, Httex1-43Q-oM8/AcK6, and unmodified mHttex1 monitored by UPLC sedimentation assay to determine the proportion of remaining monomer. **(B)** Secondary structure analysis of unmodified Httex1-43Q and Httex1-43Q-oxM8 by CD at 0 h and 48 h of aggregation. **(C)** Electron microscopy after 48 h of aggregation (scale bars = 200 nm). **(D)** Secondary structure analysis of Httex1-43Q-oxM8 and Httex1-43Q-oxM8 by CD at 0 h and 48 h post-aggregation initiation.

Interestingly, we consistently observed the presence of a population of oligomers for the oxidized Httex1 at early time points (Figure 2-C and S3). These oligomer populations were observed in a higher amount after 4 h of aggregation by EM, at which unmodified mHttex1 already showed fibrils formation (Figure S3). After 6 h, mHttex1 oxM8 showed a mixture of oligomers and short fibrils (Figure S3), whereas only fibrils were observed in the case of the unmodified mHttex1. To further validate our observations and quantitatively assess differences in dimensions and morphological properties of the aggregates formed by the two proteins, we performed time-dependent atomic force microscopy (AFM) studies. Consistent with our EM analysis, unmodified mHttex1 and mHttex1 oxM8 showed the presence of both fibrils and oligomers at 25 min, as shown in Figure 3-A. Interestingly, at 2 h, mHttex1 predominantly formed longer fibrils with a mean length of 167 nm compared to mHttex1 oxM8, which formed a mixture of oligomers and short fibrils (111 nm). Similarly, after 6 h of incubation, unmodified mHttex1 formed fibrils with a mean length of 215 nm and a height of 5.5 nm (Figure 3), whereas mHttex1 oxM8 samples showed a mixture of shorter fibrils and protofibril-like structures with average lengths and heights of 140 nm and 4.8 nm, respectively (Figure 3). AFM analysis clearly shows the effect of oxM8 on slowing the rate of fibril elongation and aggregation of mHttex1. These observations are consistent with the slower aggregation kinetics observed for the mHttex1 oxM8 in the sedimentation assay. Taken together, these results suggest that the delay in the aggregation of the oxidized mutant Httex1 could be associated with the formation and accumulation of oligomers (on- or off-pathway), that eventually convert into fibrils at later stages.

**Figure 3.**
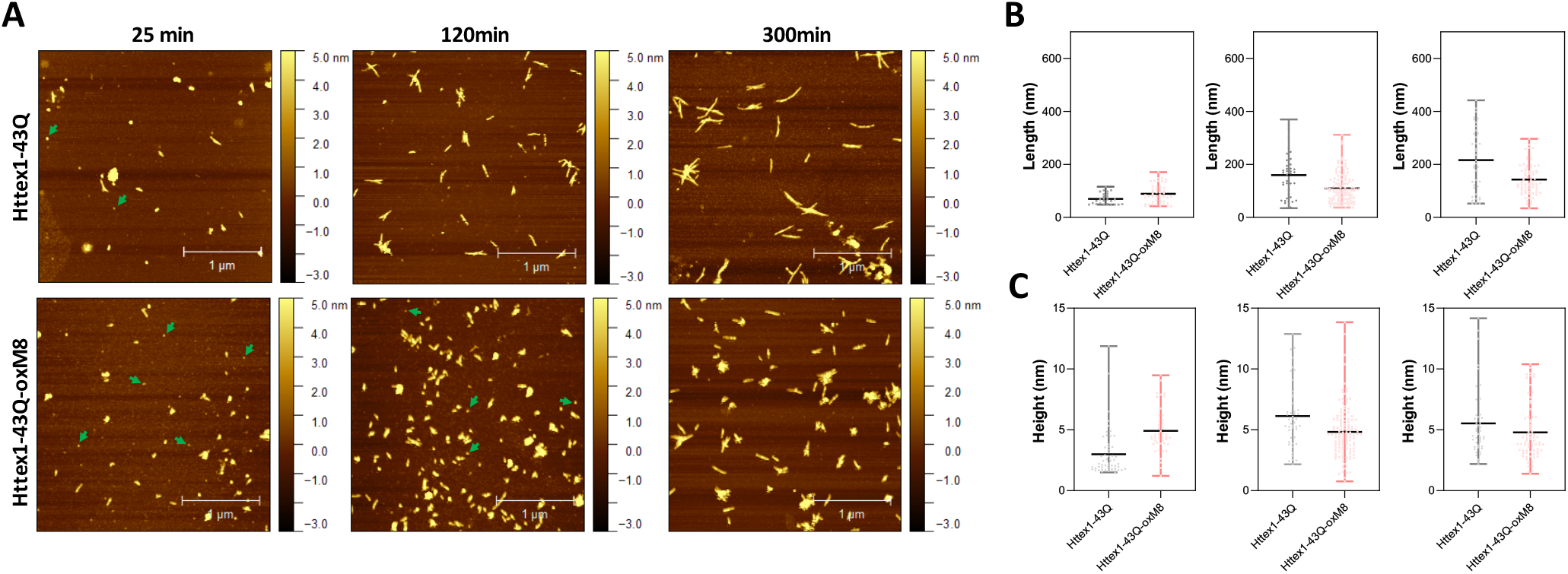
AFM analysis comparison between unmodified and Httex1-43Q-OxM8. **(A)** Time-dependent AFM images of aggregates formed by Httex1-43Q and Httex1-43Q-oxM8 (nm, nanometer), the oligomers are indicated with green arrows. Quantitative analysis of the length **(B)** and height **(C)** of fibrils measured from AFM images. All the plots for the length and height of the Httex1-43Q-oxM8 fibrils differ from the Httex1-43Q with a statistical significance of at least p < 0.05 (scattered plot represents the mean and range).

### Crosstalk between M8 oxidation and phosphorylation at T3 or Acetylation at K6

To investigate the effect of PTM crosstalk on aggregation within the Nt17 domain, we investigated and compared the aggregation kinetics and properties of Httex1-oxM8/pT3 and -oxM8/AcK6. Previously, we showed that phosphorylation at T3 significantly inhibits the aggregation of mutant Httex1 *in vitro* [30]. As shown in Figure 2, the oxidation of M8 does not seem to influence the inhibitory effect of phosphorylation at T3. Even after 48 h of incubation, ~50 % of the protein remained soluble, whereas more than 98% of the unmodified proteins have been converted to fibrillar aggregates (Figure 2-A). The CD spectrum showed that the remaining protein retains a random coil signature, suggesting that the remaining soluble protein represents monomers or disordered soluble oligomers (Figure 2-D). Analysis by EM showed the accumulation of oligomers and short fibrils similar to those we previously reported for mHttex1-pT3 [30]. On the other hand, oxidation of M8 in the context of the K6 acetylated mutant Httex1 dramatically altered the aggregation properties of the protein. Previously, we have shown that acetylation of K6 or K9 does not significantly influence the aggregation of mutant Httex1. However, as shown in Figure 2-A, the presence of both acetylation at K6 and methionine oxidation at M8 (mHttex1-oxM8/AcK6) results in significant inhibition of mutant Httex1 aggregation. Only 8% of the mHttex1 oxM8/AcK6 aggregated after 48 h (Figure 2-A) as determined by the sedimentation assay and confirmed by CD, EM, and AFM, which show predominantly disordered conformation and the accumulation of only oligomers and small fibrils (Figure 2-C and Figure 3). These results suggest that the addition of oxM8 reversed the aggregation of Ack6 and illustrates how the combination of different Nt17 PTMs could differentially influence the aggregation properties of the protein.

### Oxidation at M8 decreases the helicity of the Nt17 peptides

To gain insight into the mechanisms by which oxM8 and its combination with other PTMs influence the aggregation of mutant Httex1, we sought to investigate how these individual PTMs and their combined influence on the structural properties of the Nt17 domain. The CD spectrum of the Nt17-WT peptide showed a minimum at 200–205 nm and shoulder around 222 nm (Figure S4, Table S1) with a calculated helical content of 11%. This is in accordance with previous observation suggesting that the Nt17 domain is predominantly disordered with a tendency to form transient α-helical structures due to its amphipathic nature [29, 30, 53]. We confirmed that Nt17-AcK6 exhibited increased helical content (17%) [54] compared to the Nt17-WT peptide (Figure S4), as previously reported [54]. Additionally, by CD, we observed that Nt17-pT3 has a higher propensity to adopt helical conformation compared to other Nt17 peptides, with a helical content of 32% (Figure S4, Table S1). In contrast, when oxM8 was added to the three peptides, we observed a reduced CD signal at 222 nm for Nt17-oxM8, Nt17-oxM8/pT3, and Nt17-oxM8/AcK6 (Figure S4), indicating a decrease in the helical content which was confirmed by a fit to secondary structure content (See Table S1).

Furthermore, to investigate if the decrease in the helicity induced by oxM8 persists even at a high concentration of the peptides, we conducted CD studies (Figure S5-A) at different Nt17 peptide concentrations (60, 90, 120, 150, and 200 μM). As shown in Figure S5, all of the peptides showed a small increase in the helical content with increasing concentrations (Figure S5-B). oxM8 slightly decreased the helical content of the WT and Nt17-pT3 peptides over the different concentrations, and the difference in helicity with or without oxidation was consistent at each concentration. However, the effect of oxM8 was stronger when it was added to the AcK6 Nt17 peptide; it dramatically decreased its helical content over the whole range of tested concentrations (Figure S5-B). Taken together, these results suggest that the oxidation of M8 decreases the helicity of the Nt17 in the presence or absence of PTMs, and this can play a role in modulating the aggregation of mutant Httex1.

To better understand how oxM8 and other PTMs influence the conformational properties of Nt17, we conducted atomistic molecular dynamics (MD) studies on Nt17. For these studies, we decided to include the first two glutamine residue as preliminary studies suggested that they could be involved in stabilizing Nt17 conformations. Previously, it has been shown that there is no difference in the overall structure between the Nt17 and the Nt19 (Nt17 + QQ)[30]. We simulated the Nt19 with single and double PTMs to analyze the crosstalk between oxidation at M8, and phosphorylation at T3 or S13, or acetylation at K6. The simulations were run for a total of 13 μs using the CHARMM36m force field and a modified TIP3P water model [55], which have been shown to be effective for the simulation of intrinsically disordered proteins. The phosphate group of phosphorylated threonine was in a dianionic form. The parameters for phosphorylated threonine and acetylated lysine were obtained from the CHARMM36 parameters for modified residues (see Methods).

Both CD and MD showed that oxidation at M8 decreased the overall helicity of Nt19 unmodified, Nt19-pT3, and Nt19-AcK6 (Table 1). The CD experimental data showed that for unmodified Nt19 and Nt19-pT3, oxidation at M8 resulted in a decrease of overall helicity by 18% and 6%, while a more significant effect was observed for Nt19-AcK6, whose helicity decreased by 29% (Table 1). Likewise, the MD results also showed that oxidation at M8 decreased the overall helicity of Nt19, with a more significant decrease in helicity for Nt19-AcK6 (Table 1). The residue-wise helicity calculated from MD showed that the core region of Nt19 (residue 4-13) had a more significant decrease, while the N-term residues (1-3) were less affected (Figure 4). Overall, oxM8 has a higher impact on Nt19-AcK6 and on the core region of Nt19 in general. These observations are consistent with our aggregation studies and CD analysis data.

**Figure 4.**
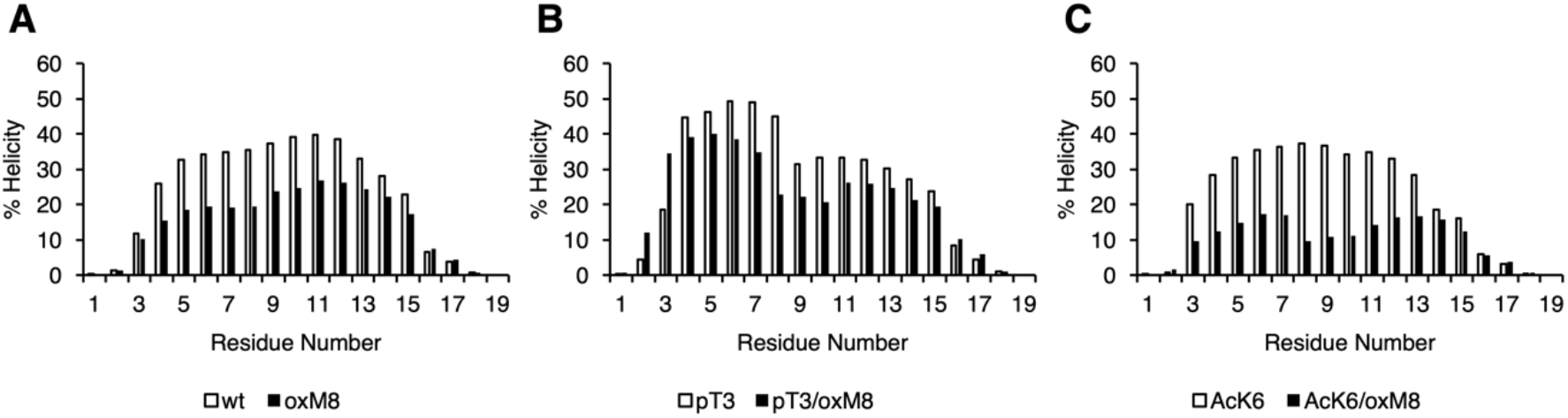
Helical propensity of Nt19. The secondary structure was obtained with DSSP from the MD trajectories of (A) WT and oxM8 (B) pT3 and pT3/oxM8 (C) AcK6 and AcK6/oxM8. The helicity for each residue was calculated based on the sum of the total number of frames that are in α-helix or 3_10_ helical conformations divided by the total number of frames.

**Table 1.** Helicity of Nt19. CD and MD both showed decrease in helicity when M8 was oxidized.

| **Nt19** | **WT vs. oxM8** | **pT3 vs. oxM8/pT3** | **AcK6 vs. oxM8/AcK6** |
| --- | --- | --- | --- |
| **MD**<br>% change in helicity | -20 | -21 | -60 |
| **CD**<br>% change in helicity | -18 | -6 | -29 |

### The formation of a short N-terminal helix in the simulations correlates with suppressed aggregation *in vitro*

The MD results were further analyzed to explore the structure and dynamics of PTM crosstalk to gain insight into the structural basis underlying the differential effects of Nt17 PTMs on the aggregation of mutant Httex1. We first performed a principal components analysis of the backbone dihedral angles (dPCA) [56] to distinguish among the various conformations adopted by the peptides with different PTMs. The first two components that depict a large part of the differences in the conformations were used for generating an energy map (Figure 5). dPCA showed well-separated minima corresponding to various helical forms and revealed a very broad and shallow minimum corresponding to a wide-range of unfolded conformations connected by very low barriers.

**Figure 5.**
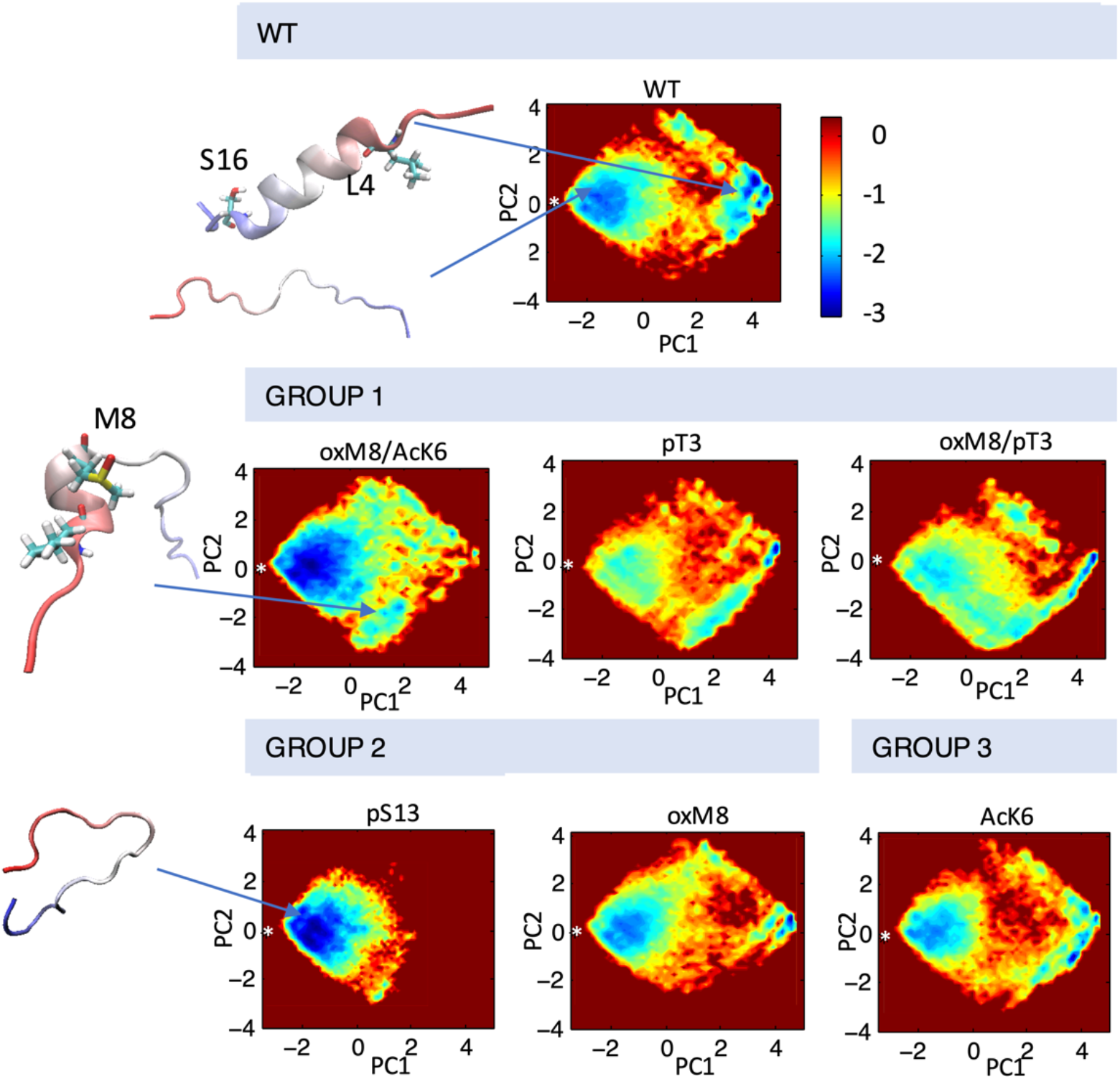
Conformational ensembles of Nt19 with different PTMs. dPCA was performed on dihedral angles calculated for the Nt19 backbone, and a free energy map was generated with the first two components. The white asterisk indicated the starting structure (in extended conformation). The major structures that correspond to local minima include fully disordered conformation, short N-term helix, and long helix. The peptides are classified based on the effect of each PTM on helicity from CD and aggregation rate: oxM8/Ack6, pT3, and oxM8/pT3 increased Nt19 helicity and decreased aggregation rate (GROUP 1), pS13 and oxM8 decreased Nt19 helicity and decreased aggregation rate (GROUP 2), AcK6 had no significant effect on aggregation rate (GROUP 3).

To investigate the connection between the major conformations and the aggregation rates of Htt with different PTMs, we classified them into three groups based on the aggregation data and overall helicity from CD data. In Group 1, we grouped PTMs that showed higher helicity in CD data and had a lower aggregation rate. In Group 2, PTMs showed lower helicity in CD data but also a lower aggregation rate, while in Group 3, PTMs showed an aggregation rate similar to unmodified Httex1. dPCA showed that for all PTMs in Group 1, there were several local minima corresponding to conformations where the first 8 N-terminal residues adopted a helical conformation while the rest remained disordered (Figure 5). When T3 was phosphorylated, and when both T3 and OxM8 modifications were present, this short N-terminal helix was 7-times more populated than for the unmodified protein. Meanwhile, the N-terminal helix was 4-times more populated in oxM8/AcK6 compared to the unmodified peptide, yet the abundance of all other helical forms was reduced by half compared to the unmodified peptide (Figure 6). For Group 2, the abundance of the N-terminal helix of pS13 and oxM8 was similar to the unmodified peptide, but a 94% and 30% decrease was observed for other types of helices in pS13 and oxM8. For Group 3, the abundance of either short N terminal helix or other types of helices in AcK6 was similar to that of the unmodified peptide. By comparing the helicity data with aggregation data, it then appears that the higher abundance of a short N-terminal helix and lower abundance of other helical conformations, like a long helix or short C-terminal helix in the MD simulations, correlates with lower aggregation rates.

**Figure 6.**
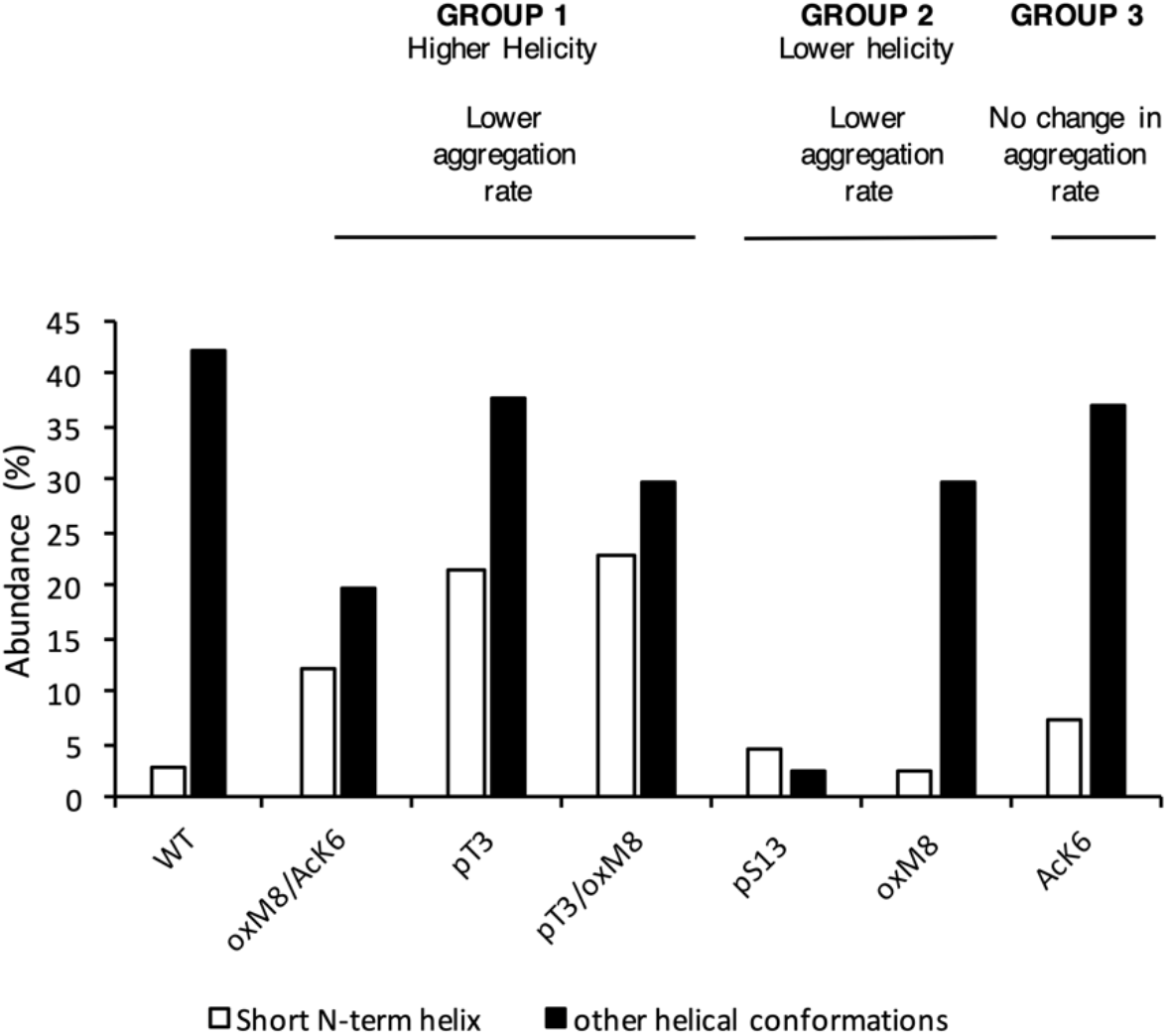
The abundance of N-term helix was high for PTMs that delayed aggregation. The abundance of short N-terminal helix and other helical forms was calculated for each peptide. Peptides with a lower aggregation rate showed a higher abundance of N-terminal helix and a lower abundance of other helical conformations.

### Oxidation of M8 increased abundance of the short N-terminal helix

To analyze how the PTMs modulate the stability of the short N-terminal helix, we searched for the key differences in structural dynamics in simulations of peptides with oxidized and reduced methionine. For this, we computed the distance matrix between Cα atoms for all frames of each MD simulation (Figure S6), compared the distributions of Cα pairwise distances through Kullback-Leibler divergences of two distance matrices, as described in Methods, and plotted the divergences into heat maps. Heat maps comparing pT3 with WT and oxM8/pT3 with WT (Figure S6) show a significant change in the distance between the first eight residues and residue 1 or 2. Heat maps comparing WT and oxM8/AcK6 show that significant changes occur at a distance between residue 4 and residue 9/10.

The distance distributions (Figure 7-A) are similar for pT3 and pT3/oxM8 but were different for WT. The peak for oxM8/pT3 and pT3 corresponds to a conformation where the phosphate group of pT3 forms a salt bridge with the NH_3_^+^ group of M1 (Figure 7-C), which is not observed in WT, where M1 and T3 remained apart. A previous study by Yalinca et al. has shown that phosphorylation stabilizes helical conformation between A2 and L7, because of the increased negative charge that counterbalances the helix dipole [57]. Also, the distribution of distances between Cα of L4 and K9 in WT, AcK6, and oxM8/AcK6 (Figure 7-B) is similar for WT and AcK6 but different for AcK6/oxM8. The peak for WT and AcK6 corresponded to the distance between L4 and K9 when they were in a helix conformation. The peak for AcK6/oxM8 corresponds to a conformation where there was a turn at M8, so the distance between L4 and K9 decreased.

**Figure 7.**
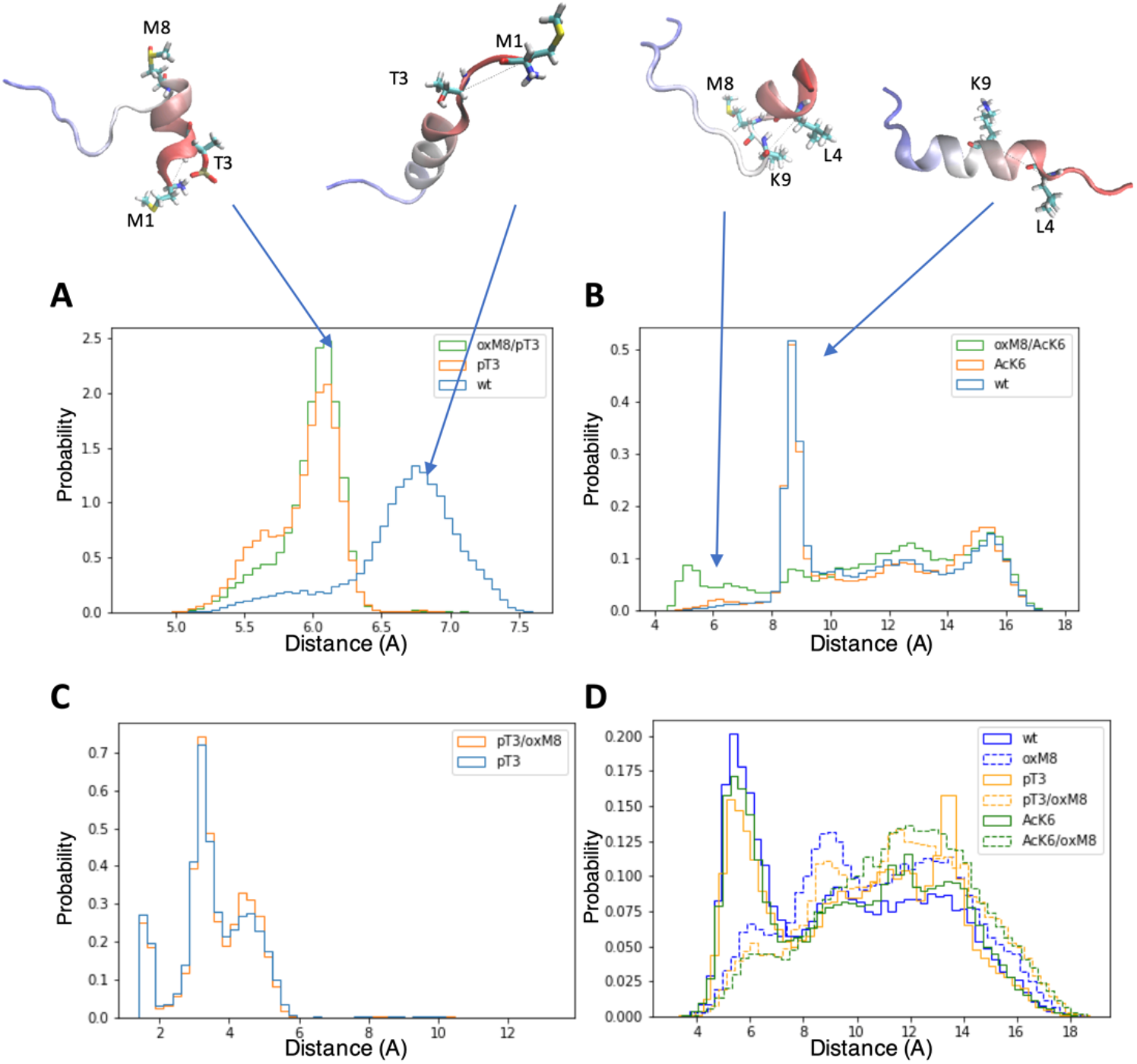
Distance distribution of atom pairs in Nt19 with different PTMs. **(A)** Distance between Cα of M1 and Cα of T3 was lower in pT3 and oxM8/pT3 than in WT. **(B)** The distance between Cα of L4 and K9 was lowered in oxM8/AcK6 compared to WT and AcK6 due to the bend at M8. **(C)** A peak below 4 Å was shown in the distribution of the distance between hydrogens of NH_3_^+^ and oxygens of the pT3 phosphate group, indicating pT3 might promote ionic interaction at this N-terminal end. **(D)** The distance between the sulfur atom of M8 and the center of the F11 aromatic ring increased in peptides with M8 oxidation, indicating the stabilization by the interaction of sulfur and the aromatic ring was weakened.

Another aspect we examined is the distance between the sulfur atom of M8 and the center of the aromatic ring of F11 because a previous study [58] has shown that when this distance is below 6 Å, an interaction between sulfur of M8 and the aromatic ring could stabilize the structure. For WT, pT3, and AcK6, a peak below 6 Å (Figure 7-D) was observed for the distance between M8’s sulfur atom and F11’s aromatic ring. When M8 is oxidized, the distance distribution shifts towards larger values, suggesting the suppression of stabilization effects brought by the interaction between the sulfur of M8 and the aromatic ring.

The residue pair distance analysis for WT, pT3, AcK6, oxM8/pT3, and oxM8/AcK6 shows that the change in abundance of short N-terminal helix and other helix resulted from (1) stabilization of N-termina helix by phosphate group of T3 (pT3, oxM8/pT3) and (2) breaking the long helix between residue 8 and 9 with oxidation at M8 (oxM8/AcK6) and destabilizing the long helix with disruption of the interaction between residue 8 and 11.

## Discussion

Among all Htt Nt17 PTMs, methionine oxidation remains the least well understood and studied. To address this knowledge gap, we first developed an efficient method to produce milligram quantities of highly purified untagged mutant Httex1 that is site-specifically oxidized at M8. To investigate the crosstalk between M8 oxidation and neighboring PTMs, we used protein semisynthetic and chemoenzymatic strategies developed by our group [30] to generate mutant Httex1 proteins bearing oxidized M8 and acetylated K6 (mHttex1-AcK6/oxM8) or phosphorylated at T3 (mHttex1-AcK6/pT3). These advances enabled us to investigate, for the first time, the role of crosstalk between these different PTMs in regulating Nt17 conformation and mutant Httex1 aggregation *in vitro*. We choose K6 and T3 because of their close proximity to M8 and the fact these are the only two PTMs that have been shown to significantly influence the helicity of Nt17 and modify the effect of neighboring PTMs on Nt17 structure and Httex1 aggregation[30].

Our results demonstrate that oxidation at M8 delayed the aggregation of mHttex1 but did not alter the morphology of the structure of the Httex1 fibrils. We hypothesized that this delay is caused by Methionine oxidation-induced formation of oligomers at the early stages of the aggregation, which undergo relatively slower conversion to fibrils. As shown in Figure 3 and highlighted in previous studies [29, 30, 50], oligomers are rarely observed during the fibrillization of the unmodified mutant Httex1 due to its high propensity to misfold and fibrillize. In contrast, our imaging studies of oxM8 aggregation consistently revealed the transient accumulation of oligomers at the early stages of the aggregation process (Figure S3).

Although acetylation at K6 was previously shown not to influence the aggregation of mutant Httex1, when this modification is combined with oxidation at M8, we observed a strong inhibitory effect on mutant Httex1 aggregation and the accumulation of mainly oligomeric and short fibrillar structures. In contrast, the combination of methionine oxidation and phosphorylation at T3 did not modify the aggregation inhibitory effect induced by pT3. Our findings are consistent with previous observations on the effect of methionine oxidation on the aggregation of other amyloid-forming proteins. For example, oxidation at M35 attenuates the aggregation amyloid-β(1–40) [59, 60], Aβ1-42, and the highly amyloidogenic Arctic Aβ1-40 variant [61]. Similarly, oxidation of methionine 109 and 112 in the prion peptide PrP106-126 inhibits its fibrillization *in vitro* [62]. Furthermore, methionine oxidation was shown to inhibits the aggregation of alpha-synuclein protein and promote the formation of stable oligomers [63]. In the case of Httex1, previous studies have shown that oxidation at M8 abolished aggregation of a model peptide consisting of Nt17 plus ten glutamine residues (Nt17Q10) [41].

To understand the structural basis underlying the effect of methionine oxidation and its crosstalk with pT3 and AcK6 on mHttex1, we performed CD and MD analysis of the unmodified Nt17 peptide and Nt17 peptides bearing pT3, or AcK6 in the presence or absence of the oxM8. We observed that in all the cases, methionine oxidation reduced the helical content of Nt17, independent of co-occurring PTMs and peptide concentration.

Consistent with our previous studies [29, 30], we observed that the effect of PTMs on the overall helicity of the Nt17 peptide did not correlate with their impact on the aggregation propensity of mutant Httex1 *in vitro*. However, it was noticed that all PTMs that experience a slower aggregation rate have a higher abundance of short helices at the first 8 N-term residues and a lower abundance of other helical conformations. According to the simulations, the higher abundance of short N-term helices in Nt17-pT3 and oxM8/pT3 could derive from direct stabilization via a salt bridge between the phosphate group at T3 and the NH_3_^+^ group of M1, while their higher abundance in Nt17-oxM8/AcK6 could result from breaking long helices with a turn between residue 9 and 10, as well as destabilizing the long helices by suppressing the interaction between residue 8 and 11 of oxM8/AcK6. The simulation results provide an interesting perspective, i.e., that the overall helicity only provides ensemble-averaged structure information; with the same overall helicity as measured by CD, some modified peptides contain a higher abundance of short N-term helix, and some contain a higher abundance of the long helix or short C-term helix forms. Previous studies suggested that the helical conformation of Nt17 is a major driver of the initiation of mutant Htt oligomerization and fibrillization. Our findings suggest Nt17 exists as an ensemble of different helical conformations, among which some might be vital for aggregation, while others, like short N-term helix, have no impact or even impair the aggregation process.

Our findings on the effect of the crosstalk between the different Nt17 PTMs highlight the complexity of the Nt17 PTM code and suggest that Htt’s normal function and aggregation are likely regulated by a complex interplay between different PTMs. A major challenge in addressing this complexity is the large number of possible PTM combinations, even within short stretches of sequences in proteins, such as Nt17. This, combined with the challenges of introducing multiple PTMs into proteins, has led to either abandoning efforts to investigate this complexity or resorting to studying protein fragments. Our findings here show that while working with peptide fragments provides useful insights, extrapolations of findings from these fragments to predict their properties in the context of full-length proteins are not straightforward (Figure 8). The work presented here represents our initial efforts in this direction and aims at exploring the extent to which this is possible using peptide systems in which the effect of PTMs can be experimentally and computationally measured and predicted. Also, our work demonstrates the promising potential of integrating experimental and computational approaches for characterizing molecular mechanisms.

**Figure 8.**
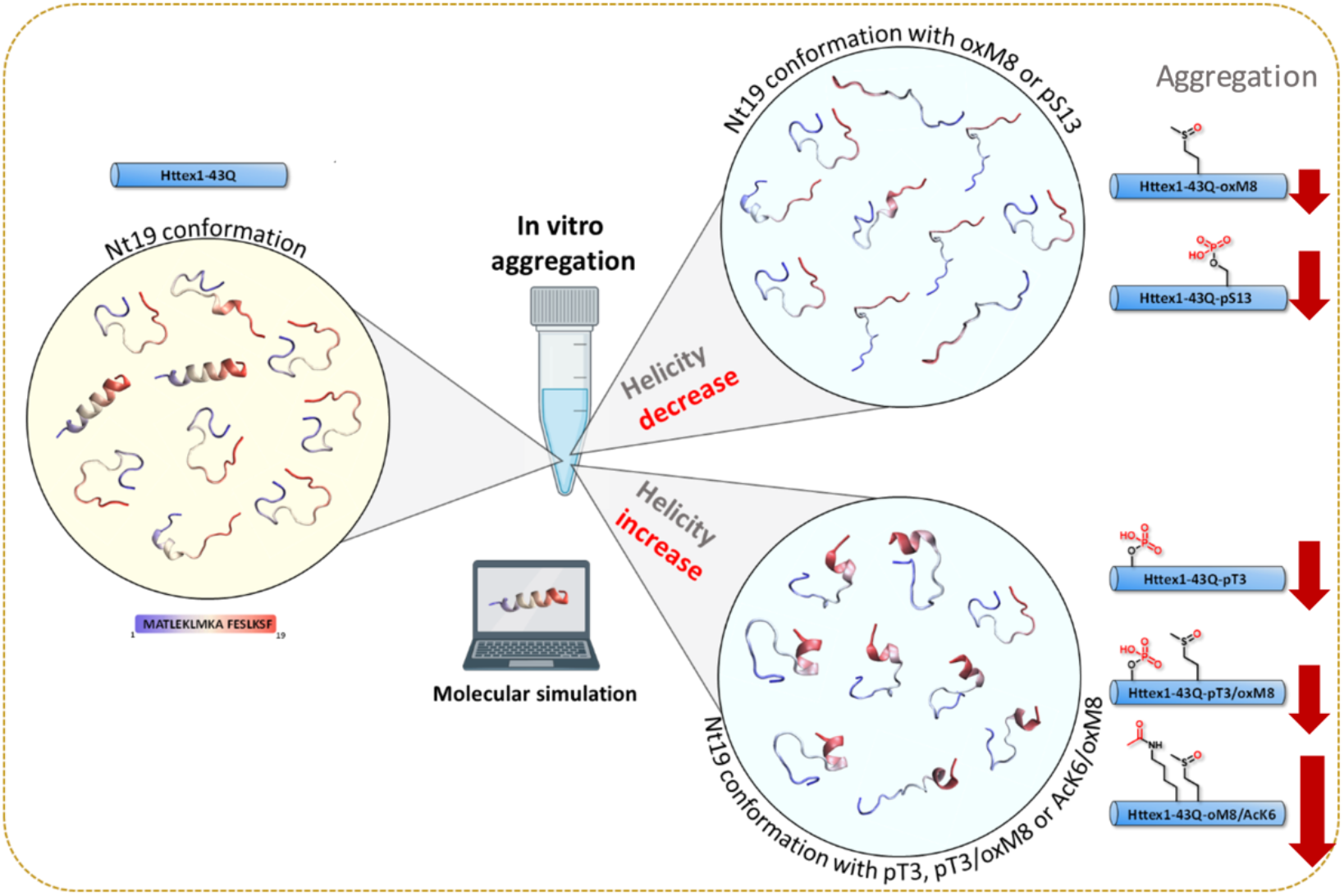
A combined experimental and computational approach unravels the structural basis of PTM crosstalk in regulating aggregation of Huntingtin exon 1.

## Materials and methods

### Materials

The pTWIN1 vector containing human Httex1 fused to His6-SUMO was ordered from GeneArt Gene Synthesis (Life Technologies); *E. coli* B ER2566 from NEB; ampicillin, DTT, isopropyl β-D-1-thiogalactopyranoside (IPTG), and hydrogen peroxide solution 30%, imidazole, complete Protease Inhibitor Cocktail, magnesium chloride (MgCl2), magnesium sulfate (MgSO4), and trifluoroacetic acid from Sigma from Sigma; PMSF from AppliChem; EGTA solution from Boston Bioproducts; Mg-ATP from Cayman; EDTA from Fisher Scientific; Luria Broth (Miller’s LB Broth) from Chemie Brunschwig; acetonitrile (HPLC-grade) from Macherey Nagel; spectrophotometer semi-micro cuvettes from Reactolab; C4 HPLC column from Phenomenex; HisPrep 16/10 column from GE Healthcare; GCK kinase (0.5 μg/ul, cat. # BC047865) from MRC PPU Reagents; Uranyl formate (UO2(CHO2)2) and Formvar/carbon 200 mesh, Cu 50 grids from EMS; high-precision cell made of Quartz SUPRASIL 1 mm light path from Hellma Analytics; Dulbecco’s Buffer Substance (PBS w/o Ca and Mg) ancienne ref. 47302 (RT) SERVA from Witech; and 100-kD Microcon fast flow filters from Merck Millipore.

### Expression and purification of SUMO-Httex1-43Q

#### Expression of His6-SUMO-Httex1-Qn (n= 23 or 43)

The expression and purification of His-SUMO-Httex1-43Q were performed as previously reported [48]. Httex1-43Q with a His-SUMO tag at its N-terminal was cloned into the pTWIN1 plasmid with ampicillin (Amp) resistance. Then, the plasmid was transformed in *E-coli* ER2566, and the resulting transformed bacterial cells were plated onto an agar plate containing ampicillin [64]. Next, 400ml LB+Amp (100 μg/ml) medium was inoculated with a single colony and incubated at 37°C overnight to start the day after the 12L expression at an optical density (OD_600_) density of 0.15. When the OD_600_ value was around 0.6, the culture was induced with 0.4 mM IPTG and incubated at 18°C overnight. The cells were harvested by centrifugation (3993 × g, 4°C, 10 min) and resuspended in buffer A (50 mM Tris, 500 mM NaCl 30 mM Imidazole, pH 7,5, 0,65um filtrated) with PMSF and protease inhibitors. The suspended bacterial pellet was subjected to sonication on ice (70% amplitude, total sonication time 5 min, intervals of 30 s sonication, 30 s pause), followed by centrifugation (39191 × g, 4°C, 60 min). The supernatant was filtered (0.45 μm, syringe filters) and passed through a Ni-NTA column. The protein was then eluted with 100% IMAC buffer B (50 mM Tris, 500 mM NaCl 15 mM Imidazole, pH 7.5, 0.65 μm filtrated). The purified fusion protein was kept ice for further reactions.

### Production of Httex1-43Q oxM8 and Httex1-43Q oxM8/pT3

SUMO-Httex1-43Q (10 mg in 18 mL) fusion protein was treated with 1 mL of 30% H2O2 to perform the oxidation directly on the SUMO fusion protein. After 2 h of the reaction, 30 μL was supplemented with 1 μl of ULP1 (1 mg/mL), and the extent of the oxidation was verified by LC/MS. 1 oxidation corresponds to +16 Da to the molecular weight of Httex1-43Q. When the completion of the oxidation was confirmed, 400 μL of ULP1 enzyme was added to the reaction solution to cleave the SUMO tag, and the reaction was directly injected in RP-HPLC, C4 column using a gradient of 25–35% solvent B (Acetonitrile + 0.1 TFA) in solvent A (H_2_O + 0.1 % TFA), over 40 min at 15 mL/min. The fractions were analyzed by LC/MS; the ones containing the protein were pooled and lyophilized. The quality of the of protein was analyzed using LC/MS, UPLC, and SDS-PAGE.

For the generation of Httex1-43Q oxM8 and Httex1-43Q oxM8/pT3, the oxidation was performed as above, and when completed, the oxidized fusion protein was dialyzed against 4 L of TBS buffer overnight at 4°C and then supplemented with 10× phosphorylation buffer to obtain a final buffer with the following composition and concentration: 50 mM Tris, 25 mM MgCl_2_, 8 mM EGTA, 4 mM EDTA, 1 mM DTT. Mg-ATP (5 mM) was added to the solution, and the pH was adjusted, and finally, GCK was added at a ratio of 1:30 w/w (kinase:protein) to the Httex1-43 oxM8 (666 μL), and the reaction was incubated at 30°C overnight. Next, the extent of the phosphorylation was verified by LC/MS, and when it was completed, 400 μL of ULP1 was added to the phosphorylation reaction solution to cleave the SUMO tag. Finally, Httex1-43Q-oxM8 was separated from the SUMO tag, and other impurities by HPLC and the fractions containing the protein were pooled and analyzed by LC/MS, UPLC, and SDS-PAGE.

### Semi-synthesis of Httex1-43Q oxM8/AcK6

The semi-synthesis of Httex1-43Q oxM8/AcK6 was performed as previously reported [30]. Briefly, 5 mg of Htt-A10C-90-43Q with free N-terminal cysteine was dissolved in neat TFA, and the TFA was dried under nitrogen after 30 min. The dried Htt-A10C-90-43Q was redissolved in 5.0 mL ligation buffer (8 M urea, 0.5 M L-Proline, 30 mM D-Trehalose, 100 mM TCEP and 100 mM MPAA). The pH was adjusted to 7.0 with 10 M NaOH, and the peptide Ac-2-9Nbz AcK6/oxM8 peptide (2.5 mg) was added to the reaction mixture. The native chemical ligation was monitored by LC-ESI-MS until the complete consumption of A10C-90 43Q. Next, the ligation reaction was desalted with hitrap 26/10 desalting column, and fractions containing the Httex1-A10C-43Q oxM8/pT3 were pooled and lyophilized. To recover the native alanine, the protein mixture was disaggregated and dissolved in 100 mM TCEP, 40 mM L-methionine, 20 vol% acetic acid in H_2_O. Freshly prepared nickel boride suspension (1.0 mL) was added to the resulting solution, and the resulting suspension was incubated at 37°C. After 2 h, a loss of 32 Da was observed. Insoluble nickel was removed from the reaction mixture via sedimentation (4°C, 4000 g, 10 min). The supernatant was subjected to RP-HPLC purification with a gradient of 25–55% buffer B (MeCN + 0.1% TFA) in buffer A (H_2_O + 0.1% TFA). The fractions containing Ac-Httex1-43Q oxM8/pT3 were analyzed by LC-MS and UPLC, pooled, and lyophilized. The final purity of the protein was checked by LC/MS, UPLC, and SDS-PAGE.

### UPLC-based sedimentation assay

To perform aggregation studies, Httex1 was disaggregated as previously reported [48] using TFA, and after evaporating the TFA, the protein was filtered through a 100-kDa filter. The final volume was adjusted with PBS to have the desired concentration and incubated at 37°C. To determine the percentage of the remaining soluble monomer during the aggregation, 40 μL was taken at each indicated time point, and insoluble aggregates were removed by centrifugation (4°C, 20000 g, 30 min). The supernatant was injected into the UPLC. The peak area was measured for each time point, and the changes in the area were used to calculate the fraction of soluble protein compared to t=0 being 100%. The exponential decay function *y* = *Ae*^*b*(*x*)^ + *c* was used to fit the data of different aggregation curves. All the mutant Httex1 aggregated protein curves had an adjusted R^2^ ranging from 0.97 to 0.99 (except Httex1-43Q AcK6/oxM8 with R^2^ = 0.78)

### Electron microscopy

For TEM analysis, 5 μl of aggregation solution was spotted onto a Formvar/carbon-coated 200-mesh glow-discharged copper grid for 1 min. The grid was then washed 3× with water and stained for 3×10 s with 0.7% w/v uranyl formate. Imaging was performed on a Tecnai Spirit BioTWIN electron microscope equipped with a LaB6 gun and a 4K × 4K FEI Eagle CCD camera (FEI) and operated at 80kV.

### Atomic force microscopy (AFM) imaging

AFM was performed on freshly cleaved mica discs that are positively functionalized with 1% (3-aminopropyl)triethoxysilane (APTS) in an aqueous solution for 3 min at room temperature. For fibril deposition on the substrates, a 20-μL aliquot of 5.2 μM protein solution was loaded on the surface of mica discs at defined time-points. Deposition took 3 min and was followed by a gentle drying by nitrogen flow. The mica discs were stored in a desiccator for 1 day before imaging to avoid prolonged exposure to atmospheric moisture. AFM imaging was performed at room temperature by a Park NX10 operating in a true noncontact mode that was equipped with a super sharp tip (SSS-NCHR) Park System cantilever. The length and height quantifications were performed semi-automatically using XEI software developed by Park Systems Corp by taking advantage of the grain detection method and thresholding (masking) the background values. The quantified data was plotted and analyzed in GraphPad Prism 8.

### Circular dichroism

mHttex1 proteins (100 μL) during the aggregation or 120 μL of the Nt17 peptides (at 60 μM in PBS) were removed from the mixture and analyzed using a Jasco J-815 CD spectrometer and a 1.0 mm quartz cuvette. Ellipticity was measured from 195–250 nm at 25°C. The spectra were smoothed using a binomial filter with a convolution width of 99 data points, and the resulting spectra were plotted as the mean residue molar ellipticity (θMRE). Then, the helical content was calculated for each peptide based on three different formula (see below), and the final reported helical content for each peptide was the average resulting value:

formula 1 [65]: 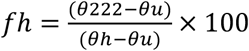, where θ222 represents the CD value at 222 nm in mean residue molar ellipticity, *θu* = −3,000 and *θh* = −39,000
Formula 2 [66]: 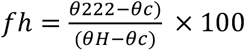, where θ222 represents the CD value at 222 nm in mean residue molar ellipticity, *θc* = 2220 – 53*T* and 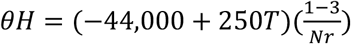 (T is the temperature, and Nr is the number of residues).
Formula 3 [67]: 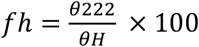, where θ222 represents the CD value at 222 nm in mean residue molar ellipticity, 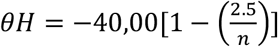 and n = number of residues.

### Molecular dynamics simulations

The starting structure for molecular dynamics simulations was a fully extended form of Htt 19 peptide. The length of this extended conformation was 68 Å, and the buffer distance between each side of the peptide and the periodic boundary was 10 Å, resulting in a water box with a width of 88 Å. The HTT 19 peptide was solvated using explicit CHARMM36m [68] modified TIP3P water model [55] and 20 mM K^+^ and Cl^-^ ions. The simulation was conducted with GROMACS [69], and CHARMM-GUI was used to generate inputs. A CHARMM36m forcefield was used with modified residues for phosphorylated threonine and acetylated lysine residues [70]. The phosphate group of phosphorylated threonine was in a dianionic form. The oxidized methionine used in this simulation was an R diastereomer, and its parameters were obtained from previously published papers [71]. The system was minimized with the steepest descent and equilibrated to 1 atm and 303.15 K with constraints on the backbone. The nonbonded interaction cut-off was 12 Å. The time step was 1 fs for equilibration and 2 fs for production. The Nosé-Hoover temperature [72] coupling method was used to maintain temperature, and the isotropic Parrinello-Rahman [73, 74] method was used for pressure coupling. LINCS algorithm [75] was used for H-bond. The production was run for 13 μs for each system.

### Data analyses

To perform dPCA [56], the dihedral angles were calculated for the Nt19 backbone, and a transformation from the space of dihedral angles (φ_n_, ψ_n_) to metric coordinate space was done by taking trigonometric function (sin φ_n_, cos φ_n_). Then, the transformed dihedral data of each post-translationally modified Nt19 was combined, and a PCA analysis was done. A free energy map was generated with the first two components. The secondary structure was obtained with the DSSP [76] algorithm implemented in VMD [77].

We analyzed and compared the effects of each post-translational modification of Nt1-19 on the dynamics using a statistical analysis of the distribution of pairwise distances between residues. For each MD of Nt1-19, we reduce the ensemble of conformations to a statistical description of the distances between all pairs of Cα in Nt1-19. For each pair of residues within Nt1–19, we compute the distribution of the Euclidean distances between the two Cα with a bin resolution of 0.5 Å. We, therefore, describe every MD with a corresponding distogram [78] summarizing the dynamics of the structure. The differences in residue-residue dynamics between MDs can be quantified using the Kullback-Leibler divergence [79] from a reference distogram.

