## Supplementary Data for "Investigating crosstalk among PTMs provides novel insight into the structural basis underlying the differential effects of Nt17 PTMs on mutant Httex1 aggregation"

### *Supplementary Material*

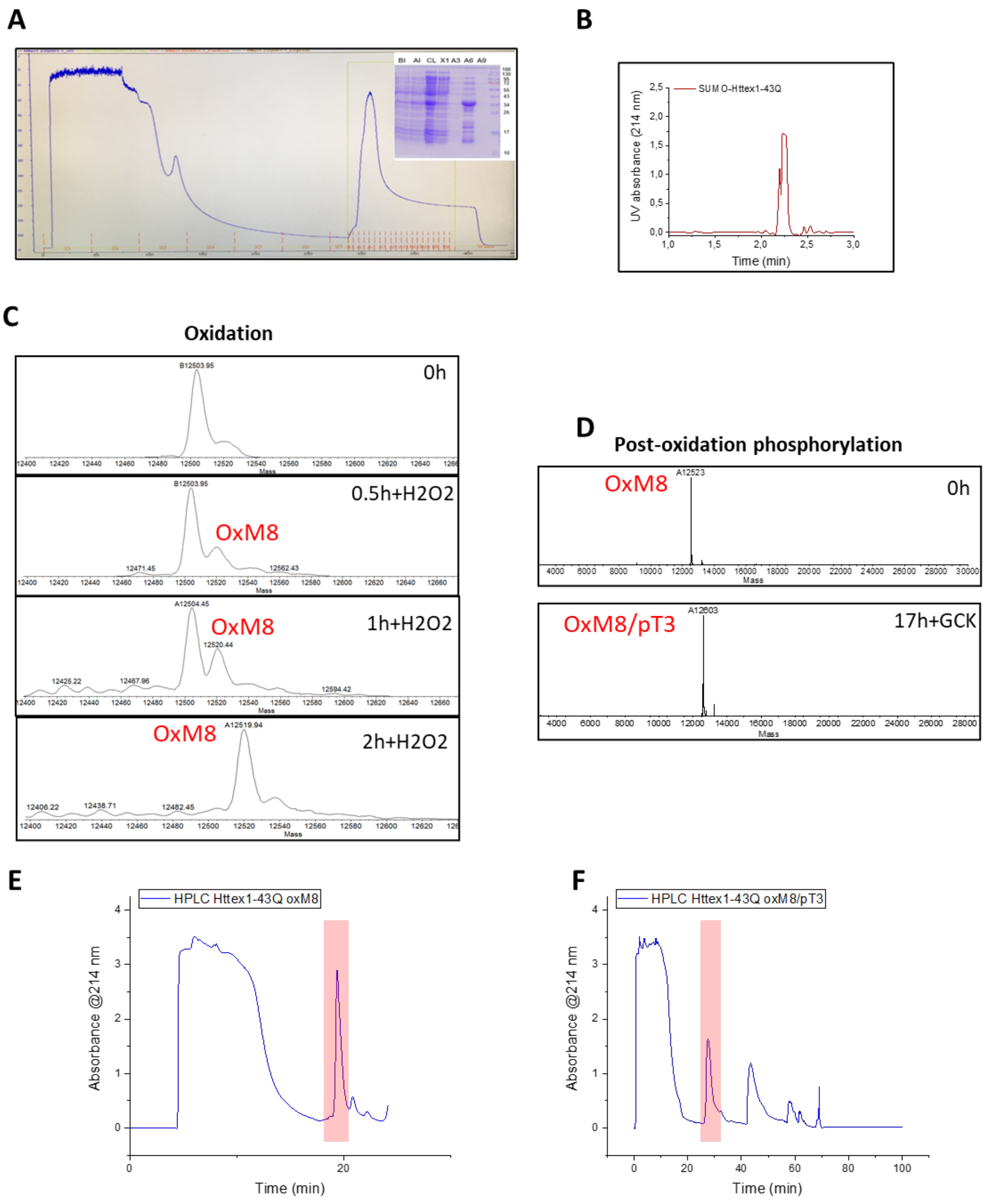


**Figure S1.** **(A)** Representative chromatogram of the IMAC purification of SUMO-mHttex1 and the analysis by SDS-PAGE of the purification fractions. **(B)** Analysis by UPLC of the fusion SUMO-mHttex1 after IMAC purifications. **(C)** Monitoring by ESI/MS of SUMO-mHttex1 oxidation by $H_2O_2$ overtime after an analytical SUMO tag cleavage by ULP1. **(D)** Monitoring of SUMO-mHttex1 phosphorylation by GCK after an analytical SUMO tag cleavage by ULP1. **(E-F)** RP-HPLC chromatograms for the purification of mHttex1 oxM8 (E) and mHttex1 oxM8/pT3 (F). (protein of interest is highlighted in red).

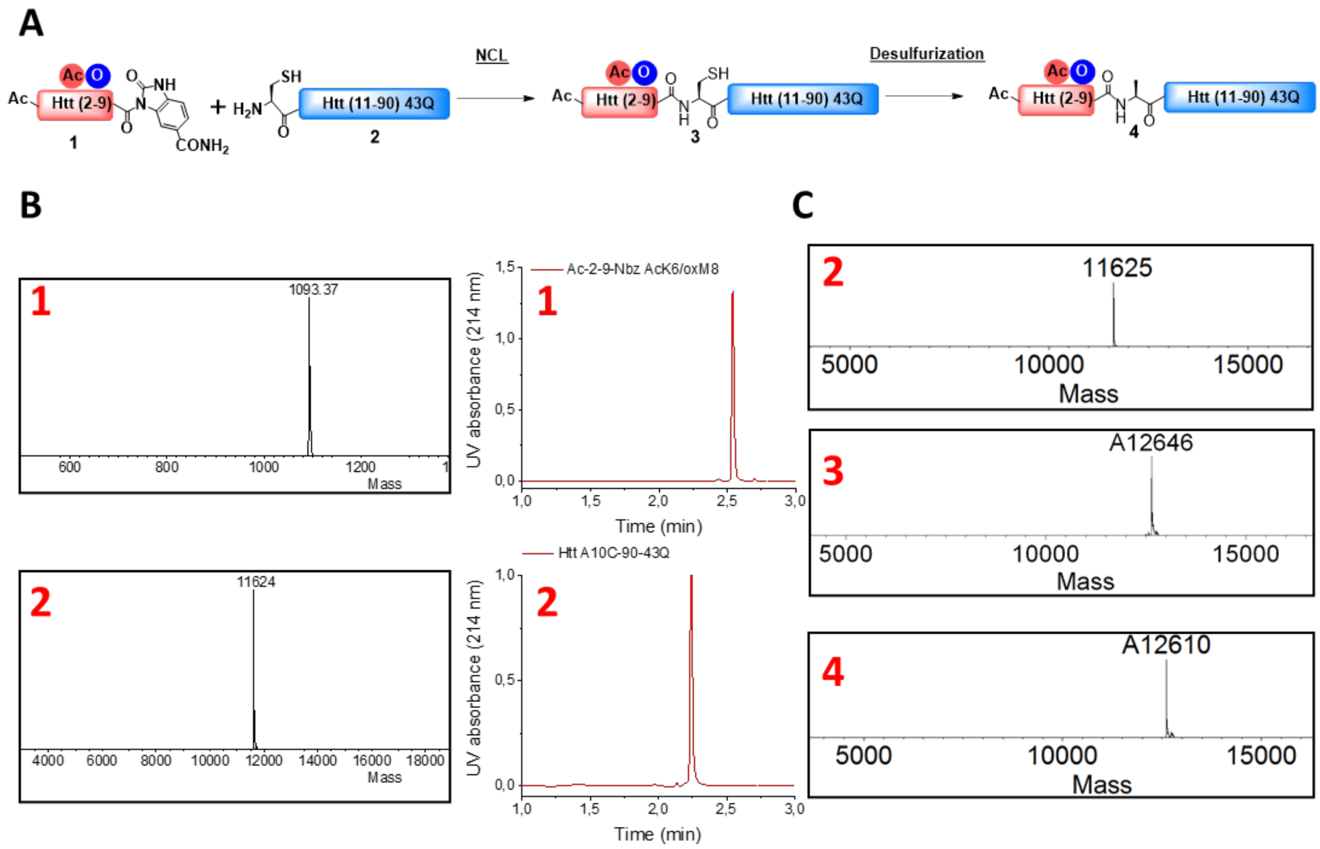


**Figure S2. (A)** Schematic representation for the semisynthetic strategy used for the generation of mHttex1-oxM8/AcK6 **(4)** (adapted from [30]). **(B)** Characterization by ESI/MS and UPLC of Htt Ac-2-9-Nbz oxM8/AcK6 **(1)** and Htt A10C-90 43Q **(2)**, both are the starting material for the semi-synthesis. The native chemical ligation of **(1)** and **(2)** was performed in (8 M urea, 0.5 M L-Proline, 30 mM D-Trehalose, 100 mM TCEP, pH 7), and the ligation was monitored by ESI/MS **(C)**. When the NCL was completed, the reaction solution containing Httex1-43Q-oxM8/AcK6 A10C **(3)** was dialyzed and lyophilized and then desulfurized in 100 mM TCEP, 40 mM L-methionine, 20 vol% acetic acid in $H_2O$ pH 1, the desulfurization of Cys to Ala was monitored by ESI/MS.

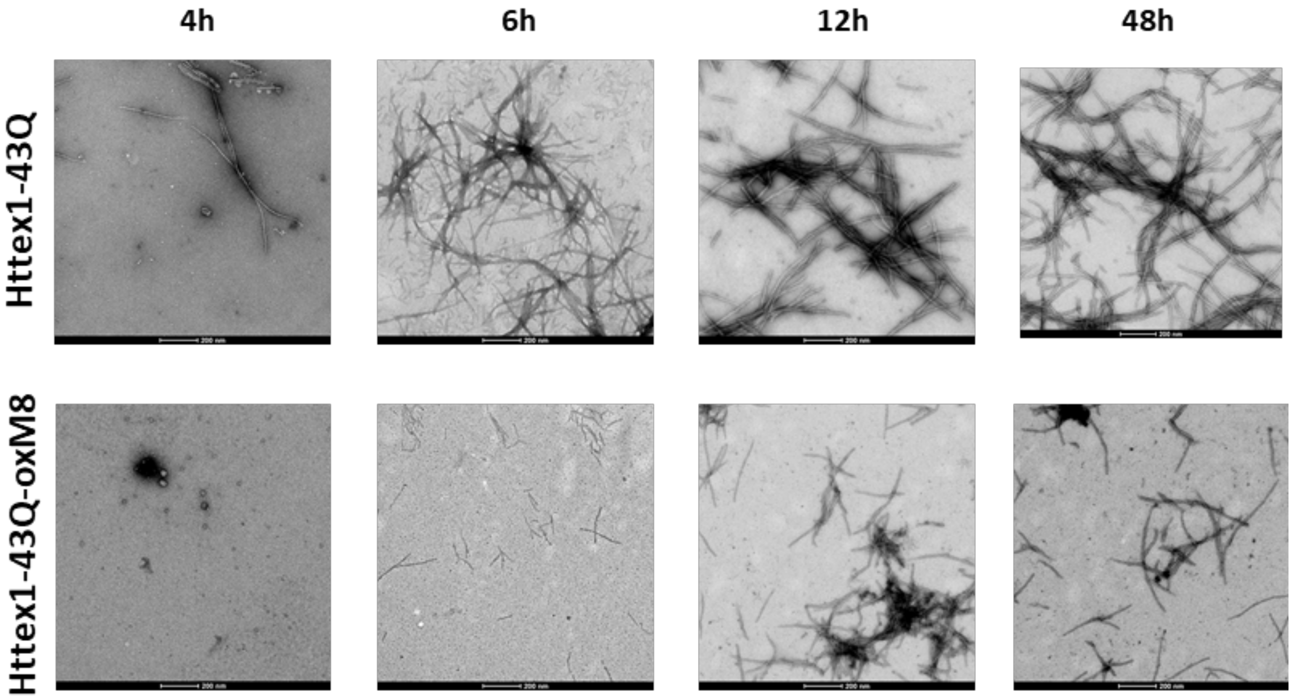


**Figure S3.** Over-time aggregation monitoring by electron microscopy of Httex1-43Q-oxM8 compared to unmodified Httex1-43Q (at 10 uM).

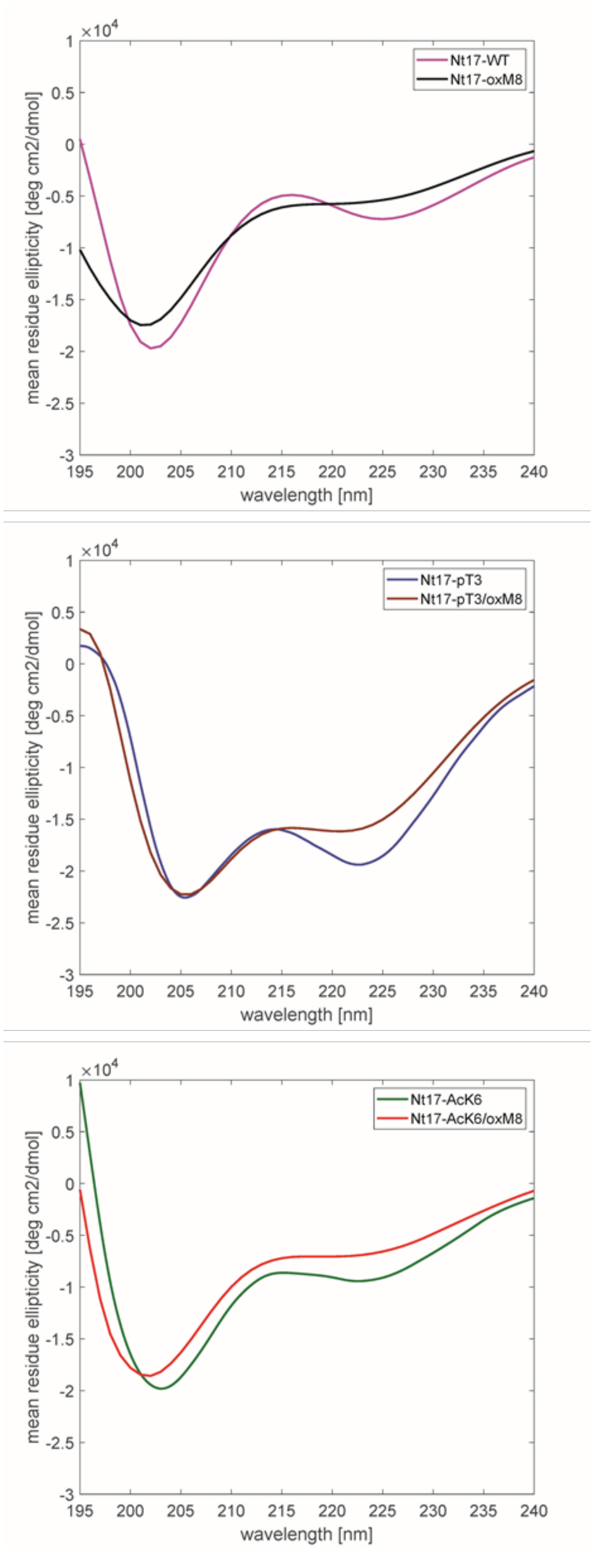


**Figure S4.** Far-UV CD spectra of Nt17-WT, and Nt17-oxM8, Nt17-AcK6, Nt17-AcK6/oxM8, Nt17-pT3 and Nt17-pT3/oxM8 at 60 μM

**Table S1:** Helical content (%) calculated for the different peptides at 60 μM.

| **Nt17 peptide** | **CD Helical content (%)** |
|---|---|
| WT | 11 |
| oxM8 | 9 |
| pT3 | 32 |
| pT3/oxM8 | 30 |
| AcK6 | 17 |
| AcK6/oxM8 | 12 |

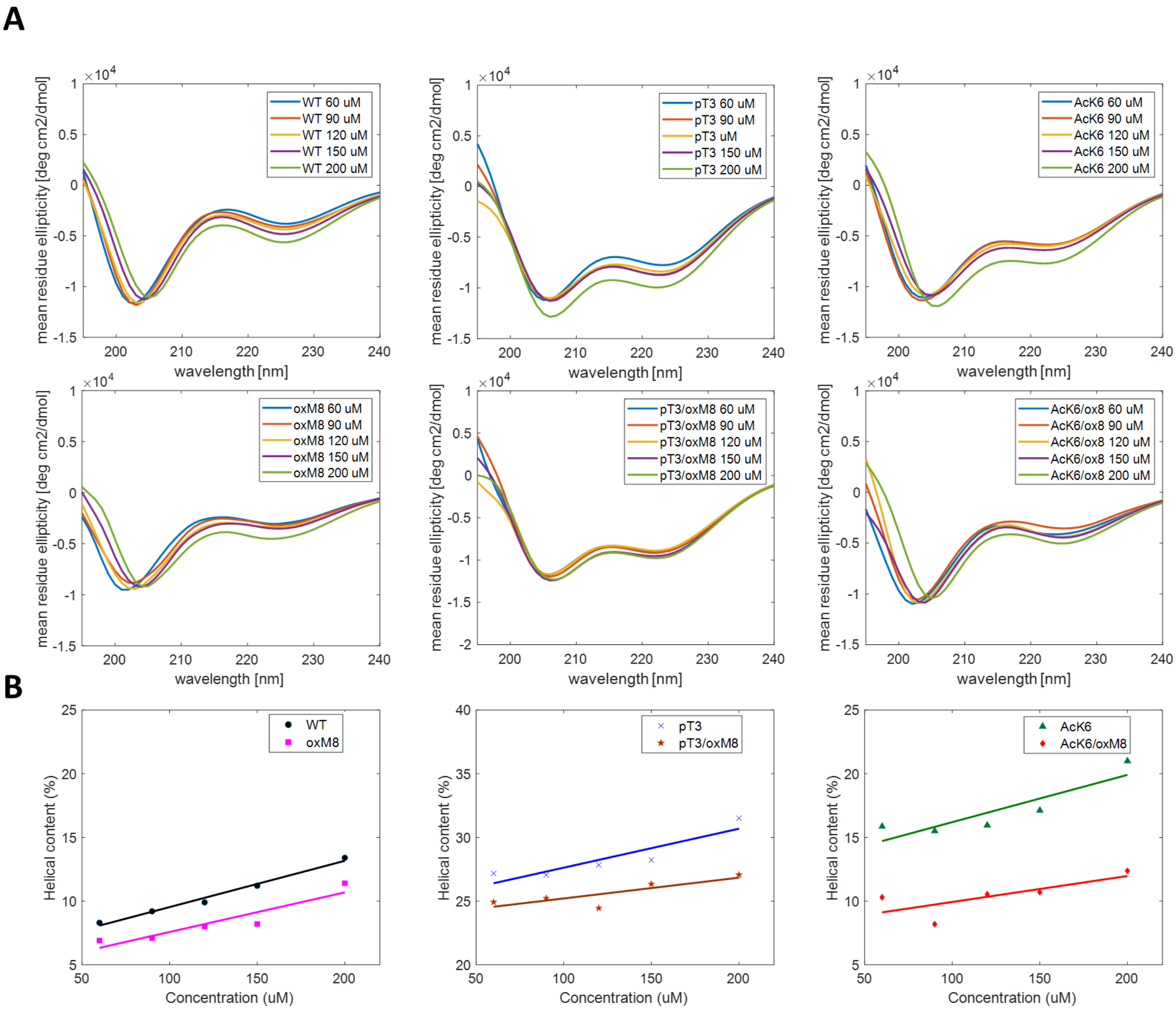


**Figure S5.** **(A)** Far-UV CD spectra of Nt17-WT, and Nt17-oxM8, Nt17-AcK6, Nt17-AcK6/oxM8, Nt17-pT3 and Nt17-pT3/oxM8 at 60, 90, 120, 150, and 200 μM. **(B)** Helical content for the peptides in the function of the different concentrations.

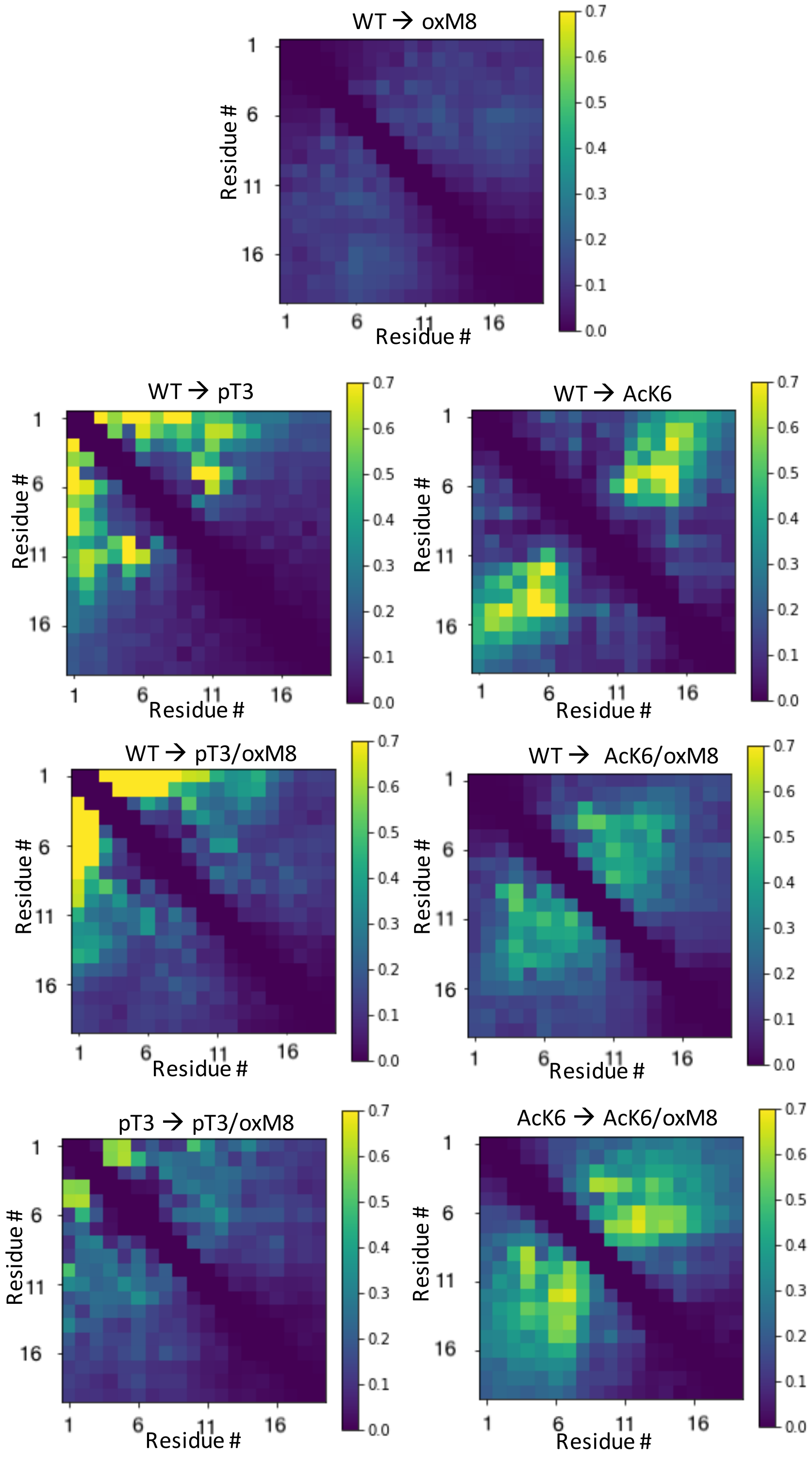


**Figure S6. The Kullback–Leibler divergence of pairwise distances of Cα.** For each MD of Nt19, the distances between all pairs of Cα in Nt19 were calculated. The differences in residue-residue contact were quantified using the Kullback-Leibler [79] divergence from its reference distogram.
